# Forestwalk: A machine learning workflow brings new insights into posture and balance in rodent beam walking

**DOI:** 10.1101/2024.04.26.590945

**Authors:** Francesca Tozzi, Yan-Ping Zhang, Ramanathan Narayanan, Damian Roquiero, Eoin C. O’Connor

## Abstract

**Abstract:** The beam walk is widely used to study coordination and balance in rodents. While the task has ethological validity, the main endpoints of ‘foot slip counts’ and ‘time to cross’ are prone to human-rater variability and offer limited sensitivity and specificity. We asked if machine learning-based methods could reveal previously hidden, but biologically relevant, insights from the task. Marker-less pose estimation, using DeepLabCut, was deployed to label 13 anatomical points on mice traversing the beam. Next, we automated classical endpoint detection, including foot slips, with high recall (>90%) and precision (>80%). A total of 395 features were engineered and a random-forest classifier deployed that, together with skeletal visualizations, could test for group differences and identify determinant features. This workflow, named Forestwalk, uncovered pharmacological treatment effects in C57BL/6J mice, revealed phenotypes in transgenic mice used to study Angelman syndrome and SLC6A1-related neurodevelopmental disorder, and will facilitate a deeper understanding of how the brain controls balance in health and disease.

## Introduction

A wide variety of tests have been developed to measure motor function in rodents (1). These include measures of gross neurological function, such as the SmithKline, Harwell, Imperial College, Royal Hospital, Phenotype Assessment (SHIRPA; (2)), to others that provide insights into fine motor skill and learning, such as directional reaching tasks (3). Motor coordination and balance is typically measured with either a rotarod, or in climbing and swimming tasks, or in beam walking and stepping tests (1,4). The beam walk was originally established to detect damage to the motor cortex in rats and mice (5–8), but has since been deployed across multiple disciplines and disease areas, including for drug pharmacology and toxicology (9,10), the study of aging (11,12), spinal cord injury (13,14), neuromuscular disorders (15), stroke (16), Alzheimer’s disease (17), Parkinson’s disease (18), Huntington’s disease (19), and multiple sclerosis (20). Of note, beam walking is also used to study balance and coordination in humans (21,22), and thus holds additional value to enable direct comparison of test performance across species.

In the beam walk, rodents are required to traverse a suspended narrow beam to reach a goal area (1,4,23). The length, shape, diameter, material and angle of the beam can be varied in order to change the complexity of the task, and reveal deficits in coordination and balance that may otherwise not be evident from kinematic observations made on a flat walking surface (24). The popularity of the test likely resides in the fact that it is simple to set-up, is inexpensive, and can be performed after limited training and with minimal expertise of the experimenter (4). The two primary endpoints typically reported in the beam walk are ‘the number of foot slips of the hind paw’, and ‘the time to cross the beam’ (1,4,18,23,25). Additional measures such as number of falls or total distance traveled on the beam may also be reported, but are typically only changed in models with more severe phenotypes (e.g. with spinal cord injury (4)).

Although widely used and easy to implement, a major limitation of the beam walk test is that its primary endpoints, including number of foot slips and time to cross, are typically scored by human-raters. Human-raters are well known to demonstrate high levels of inter- and intra-rater variability, and likely contribute to reduced reliability and reproducibility of findings. Human-rater scoring is also very time intensive (4), and is only possible with a limited number of observations that are readily identified by the human eye.

Consequently, subtle changes in behavior are challenging to quantify and can escape entirely from human detection (26–29). Together with the limited *sensitivity* of endpoints available to human-raters, a restricted set of endpoints also leads to poor *specificity* of the beam walk test to discriminate between fundamentally different experimental conditions. Numerous studies report increases in foot slips in beam walking, despite using rodent models with non-overlapping disease pathomechanisms. Taken together, achieving automated endpoint detection in the beam walk could lead to more robust and reproducible experimental findings. Furthermore, increasing both the sensitivity and specificity of beam walk endpoints to detect and discriminate subtle yet relevant functional changes is of critical importance if the test is to reliably inform on the link between brain function and posture and balance control.

Motivated by these opportunities, we set out to use modern computational neuroethology methods to gain greater insight into posture and balance in beam walking in mice (28,30–34). While other reports have also proposed alternative data capture and analysis methods for beam walking, they either still rely on human-rater scoring and annotation (13,14,35,36), or are unable to automate detection of foot slips (18,24), or have not demonstrated scalability of the proposed approach to full experimental conditions (37). To address these gaps, we first deployed DeepLabCut (DLC), an open-source toolbox that allows training of a deep neural network to annotate animal key-points to human-level accuracy (33). Using DLC key-points from each mouse recorded during beam walking, we subsequently developed an analysis workflow that fully automated detection of the classical endpoints of number of foot slips and time to cross the beam. A total of 395 features were then engineered from each beam walk test and used to train a Random Forest Classifier (RFC) to detect differences between experimental groups and reveal determinant features that, together with skeletal representations of mice, could provide greater insight into posture and balance than achieved by classical endpoints alone. Proof-of-principle for this workflow, named Forestwalk, is given by increased sensitivity to detect pharmacological effects of diazepam, and the ability to identify similarities and differences amongst transgenic mouse strains commonly used in the study of two neurodevelopmental disorders: namely Angelman syndrome and SLC6A1-related neurodevelopmental disorder.

## Results

### Forestwalk provides increased sensitivity to detect drug effects on animal posture and balance

To enable automated and detailed analysis of mouse posture in the beam walk test, we first deployed DeepLabCut (DLC), an advanced tool for markerless pose estimation (33). A deep-learning-based neural network was trained in DLC to label 13 distinct key-points on mice as they traversed the beam (**Fig. 1A**). Additionally, 5 points were labeled on the beams themselves, allowing automated identification of three different beams used throughout the study (i.e. 16 mm square, 16 mm round and 9 mm square beams, referred to as Beam 1, Beam 2 and Beam 3, respectively). These additional points also demarcated the central 80 cm section of each beam where the behavioral analysis was focused (**Fig. 1A**).

**Fig 1.**
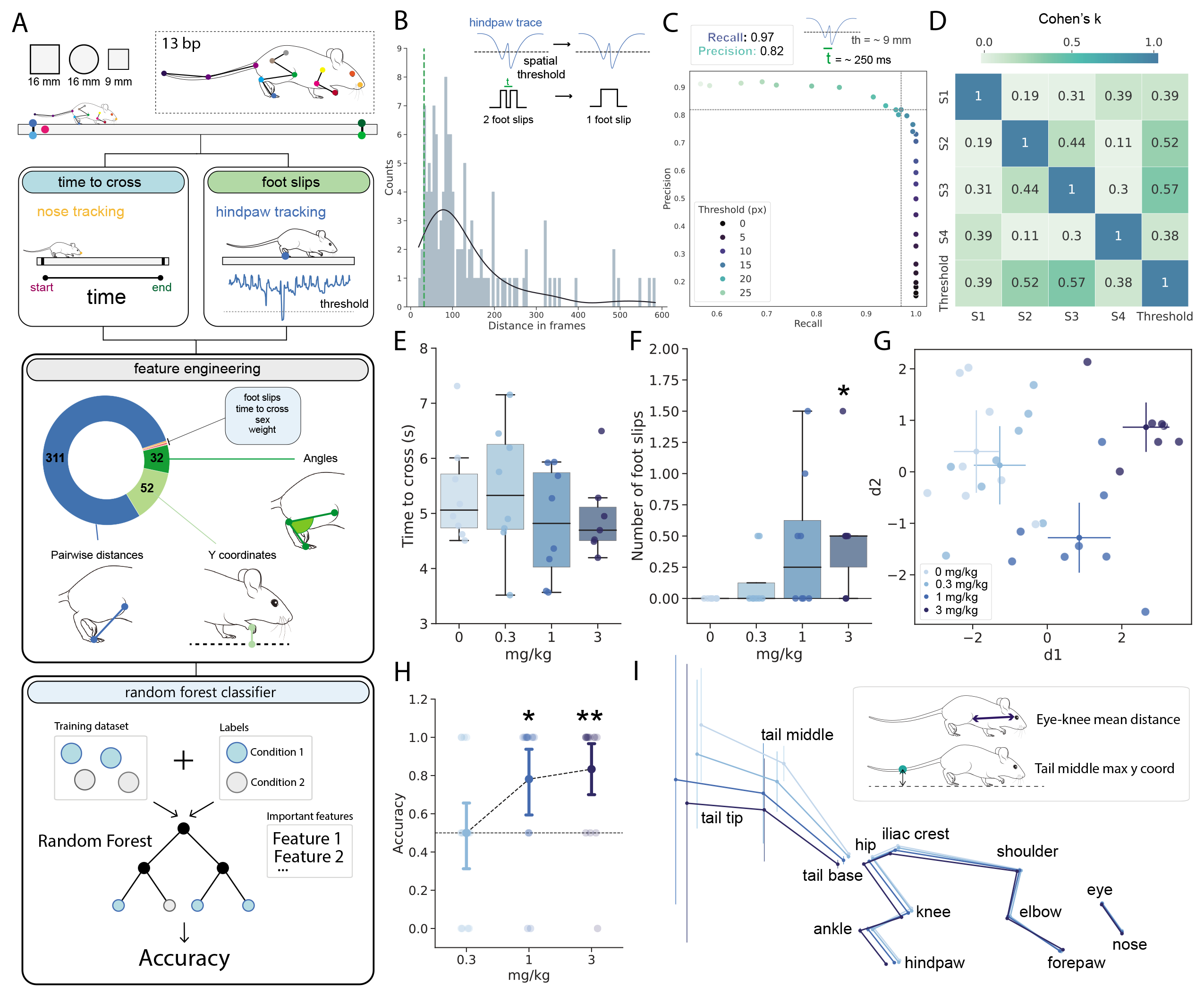
Forestwalk reveals treatment effects of diazepam in the beam walk test. **(A)** Schematic of data acquisition with DLC, and subsequent feature detection/engineering and analysis workflow (‘Forestwalk’). **(B)** Histogram with frequency of temporal gaps (i.e., frames) between foot slips (determined by experienced human-raters from 43 videos across 3 experiments). Green line marks lower 5% quantile; temporal cutoff below which consecutive slips are combined. Insert illustrates initial identification of slips using a y-axis threshold, and subsequent consolidation of two events based on temporal proximity. **(C)** Precision-Recall curve comparing performance of the threshold-based foot slip detection method, with varying thresholds, vs. experienced human-rater’s annotations on the same dataset. Dashed lines indicate the precision (0.82) and recall (0.97) achieved using the chosen y-axis threshold (18 pixels, ∼9 mm below the beam). **(D)** Concordance (Cohen’s kappa) between experienced human-raters (’S1-S4’) and the threshold-based detection method. **(E)** Time to cross the beam (Beam 2) following vehicle or diazepam administration (Effect of treatment; ANCOVA with weight as a covariate, F(3,26) = 0.86, p = 0.48). **(F)** Foot slips on the beam (Beam 2) following vehicle (i.e. 0 mg/kg) or diazepam administration. ANCOVA with weight as covariate: F(3, 26) = 2.82, p = 0.058; post-hoc analysis with Tukey’s test: p=0.05, 0 mg/kg vs. 3 mg/kg). **(G)** Two-dimensional (2D) plot with first two discriminants (d1,d2) from a Linear Discriminant Analysis (LDA) on the diazepam dataset. **(H)** Point plot indicating the classifier’s average accuracy to distinguish a specific diazepam treatment from vehicle. One-sample Wilcoxon test vs. chance level (0.5: dotted line): p = 0.023, 1 mg/kg vs chance, W = 63; p = 0.0063, 3 mg/kg vs chance, W = 65. **(I)** Skeletal representation for each group in the diazepam experiment. Points show mean position, with vertical bars showing y-axis variance for each body part. Insert schematic highlights the two most important features that discriminate between all treatment groups. For E-I: Sample size n = 8 mice. Box plots show median ± 95% CI. 2D plot of LDA shows centroid ± 95% CI. * p <0.05, ** p < 0.01.

With DLC-derived key-point data in hand, we developed an analysis workflow for the beam walk that we refer to as ‘Forestwalk’. First, we aimed to automate the recording of standard metrics most often reported in the beam walk task, namely ‘time to cross’ and ‘number of foot slips’. Such metrics are typically recorded manually by human-raters. However, this approach is prone to intra- and inter-rater variability, and likely contributes to poor reproducibility of findings from rodent behavioral neuroscience research.

The endpoint of ‘time to cross’ was obtained simply by using the position of the animal’s nose, and measuring the duration taken for the nose to enter and exit the designated central 80 cm region of the beam (**Fig. 1A**). Parts of the beam on either side of this region were excluded from this measure. Specifically, activity in the first 10 cm of the beam was subject to variability in how the experimenter placed the mouse at the start location, while locomotor behavior in the final 10 cm of the beam was often contaminated by exploratory activity directed towards the escape platform (e.g., pause in locomotion, stretch-attend posture etc).

To enable automated detection of ‘number of foot slips’, we first operationally defined a foot slip as ‘a downward deviation of the hindpaw from its anticipated trajectory along the beam’. To verify this definition, 43 videos randomly selected from three different experimental datasets (**S2 File**) were manually labeled for foot slips by an experienced human-rater. Visual comparison of manually labeled foot slips (n=162) with the hindpaw y-coordinate confirmed that human-labeled foot slips were consistently accompanied by a downward deviation in the trajectory of the hindpaw y-coordinate. Moreover, this deviation appeared to exceed the typical position of the hindpaw along the beam (see example in **Fig. 1A**). Based on this observation, we rationalized the use of a threshold-based detection method that would count a foot slip whenever the hindpaw y-coordinate fell below a defined spatial threshold. From the set of human-labeled foot slips, the average time between two consecutive slip events was 138 frames, which translates to ∼1 second at a recording speed of 120 frames per second, and the likelihood of two foot slips occurring within 32 frames of each other (i.e. ∼250 milliseconds) was less than 5% (**Fig. 1B**). Thus, to avoid automated double-counting of closely spaced foot slips, which were most likely part of one continuous foot slip event, a temporal threshold was also implemented to amalgamate any slip events that occurred within 32 frames of one another. With the temporal threshold fixed, systematic variation of the y-coordinate threshold for the hindpaw position revealed a spatial threshold of ∼18 pixels below the beam surface (equivalent to ∼9 mm) gave the highest level of recall (0.97) and precision (0.82) when compared to the experienced human-rater annotations on the same set of videos (**Fig. 1C**).

The generalizability of the threshold-based foot slip detection method was next assessed by comparisons with four experienced human-raters. The threshold-based method produced outcomes that were consistent with those of the experienced raters and demonstrated the highest levels of agreement with most of them (**Fig. 1D**). To ensure that our chosen threshold was not overly fitted to the initial dataset and experimental conditions, one experienced rater independently evaluated a new set of 18 videos sourced from different behavioral experiments (**S3 File**). Foot slip events detected by the threshold-based method were evaluated against this new, human-rater-derived ‘ground truth’. The comparison again demonstrated high levels of recall (0.92) and precision (0.81), thereby validating the method’s consistent performance and its capability to generalize across different experimental datasets. Collectively, these findings indicate that our threshold-based foot slip detection method serves as a reliable and objective standard, providing a consistent measure that closely corresponds with the judgments made by multiple experienced human scorers. By contrast, agreement between different experienced human-raters was generally poor (**Fig. 1D**), which underscores the subjective nature of manual scoring and further emphasizes the value of the automated threshold-based detection method.

With the classical endpoints of ‘time to cross’ and ‘number of foot slips’ fully automated, we expanded the capabilities of Forestwalk to take full advantage of the 13 labeled key-points. Our expectation was that deep analysis of these data points would provide more detailed insights into postural control and balance on the beam walk than afforded by use of the classical endpoints alone.

For the next part of the Forestwalk workflow a feature engineering step is first deployed. Pairwise distances between all body parts are calculated, as well as the distances from each body part to the beam surface, and essential joint angles are quantified. Each feature is initially represented as a time series, from which key statistical descriptors are extracted: the mean, minimum, maximum, and variance. This methodology, plus inclusion of body weight, sex, and the classical endpoints of time to cross and number of foot-slips, generates a dataset of 395 features for each recorded video of a mouse traversing a beam (**Fig. 1A**). To adjust for individual differences in body size that could influence the measured features, we introduced a normalization step. Distances between body parts, as well as from each body part to the beam surface, were standardized using a reference dimension - the distance between the elbow and the shoulder. This reference dimension was selected for its consistency across subjects, allowing for more precise comparisons of postural dynamics not biased by the animal’s overall body size. In addition, recognizing the significant impact of body weight on movement and balance, the weight of each animal was included as a distinct feature in the dataset and subsequent statistical analyses included body weight as a covariate.

To refine this extensive dataset and reduce multicollinearity, a feature selection step follows in the Forestwalk workflow. Recursive Feature Elimination (RFE) is deployed in tandem with a Random Forest Classifier (RFC) (38) to whittle down the feature set from 395 to the 50 most informative features that can effectively distinguish between experimental conditions. These 50 prioritized features are then used to train a RFC, which is designed to differentiate between experimental groups (e.g. drug treatment or genotypes). RFC performance is validated using a leave-one-out cross-validation scheme, assessing the predictive accuracy for individual animals. This accuracy reflects the RFC’s ability to correctly classify an animal into its true biological group (**Fig. 1A**). A classifier’s accuracy that significantly outperforms chance level would indicate a meaningful distinction between the conditions we aim to differentiate. When the average accuracy is found to be significantly higher than chance, it confirms that the conditions are distinct. A further benefit of the Forestwalk workflow is that prioritized features can be further analyzed to determine the most influential behavioral traits defining differences between experimental conditions (**Fig. 1A**).

To evaluate the effectiveness of Forestwalk in detecting changes in beam walking performance, we conducted a pharmacological study with diazepam, a benzodiazepine known for its anxiolytic properties, but also linked to side-effects of sedation and ataxia. Previous research has indicated that even low doses of diazepam can induce motor effects in humans, which are not always consistently observed in rodent models (9). Diazepam (0.3, 1 and 3 mg/kg) or vehicle (i.e., 0 mg/kg) was administered intraperitoneally to n=8 male C57BL/6J mice, 30 minutes prior to starting the beam walk test, using a within-subjects crossover design. The automated measures of ‘time to cross’ did not change with any dose of diazepam in comparison to vehicle, while the ‘number of foot slips’ significantly increased only at the highest dose of 3 mg/kg (**Fig.1 E,F** for beam 2, and note similar outcomes on other beams in **S1 Fig. A-F**). A linear discriminant analysis (LDA) applied to all 395 features indicated more nuanced treatment effects of diazepam (**Fig.1 G**). A distinct separation between animals administered with vehicle and 1 mg/kg of diazepam was already evident in LDA, which became more pronounced at 3 mg/kg, as confirmed by the increased euclidean distance from the vehicle centroid (**Fig.1 G, S1 Fig I**). To substantiate these findings, a RFC was trained to differentiate between the vehicle control and diazepam treated groups. A significant difference between vehicle and 1 mg/kg diazepam was identified, and the difference was even more pronounced between vehicle and 3 mg/kg diazepam (**Fig. 1H**). Taken together, these data highlight the superior sensitivity of Forestwalk to detect pharmacological effects of diazepam on beam walking performance, which were not evident from the two classical endpoints of ‘time to cross’ and ‘foot slips’ alone.

An average skeletal representation of mice from each treatment group was generated to further explore potential postural changes following diazepam administration (**Fig. 1I**). A progressive elongation of the body and a lowering of the center of mass with increasing doses of diazepam was observed, along with a downward shift in tail position (**Fig. 1I** and see an example video in **S4 File**). In line with these observations, the mean distance from eye-to-knee and the maximum y-coordinate of the tail’s central point were identified by Forestwalk as the top-ranked prioritized features in discriminating between the different diazepam doses (**Fig. 1I**; see **S2 File**). Thus, as the dose of diazepam increases, there is a corresponding progressive impact on the animals’ posture and tail positioning. Forestwalk is sensitive to identifying these subtle but relevant changes in body posture and balance, and enable their quantitative utilization for behavioral phenotyping.

### Forestwalk reveals previously hidden phenotypes in mouse models for Angelman Syndrome

The beam walk test is often used to study genetic models for neurological disorders, and thus we asked if Forestwalk could provide new insights in such a context. We focused first on mouse models for Angelman Syndrome (AS), a neurodevelopmental disorder marked by the absence of the maternal Ube3a protein (39,40). Individuals with AS and mouse models used for AS research are reported to exhibit impaired movement and balance. However, the extent of motor impairments varies across different AS transgenic mouse models, and the underlying causes remain elusive (41–45).

We focused on three commonly used mouse models: 1) Ube3a 129 mice (129-Ube3a^tm1Alb^/J, Jax ID: 004477), with a targeted deletion in maternal Ube3a exon 2 (46); 2) Ube3a B6/129 hybrid mice (B6.129S7-Ube3a^tm1Alb^/J, Jax ID: 016590), with a similar deletion but in a mixed genetic background (46); and 3) AS-ICTerm mice (C57BL/6J-Rr70^em1Rsnk^/Mmmh, 065423-MU), which has a disruption in the maternal imprinting center affecting Ube3a and several other genes (41). All lines were bred to obtain wild-type (WT) littermates to serve as controls to the respective Ube3a mutant animals (commonly referred to as ‘KO’). Ube3a mutant mice have previously been reported as overweight compared to WT controls (42,43). Likewise in our study, a comparison of body weight indicated significant genotype-related differences in the Ube3a 129 (**Fig. 2A**) and Ube3a B6/129 (**Fig. 2E**) strains, but not in the AS- ICTerm mice (**Fig. 2I**).

**Fig 2.**
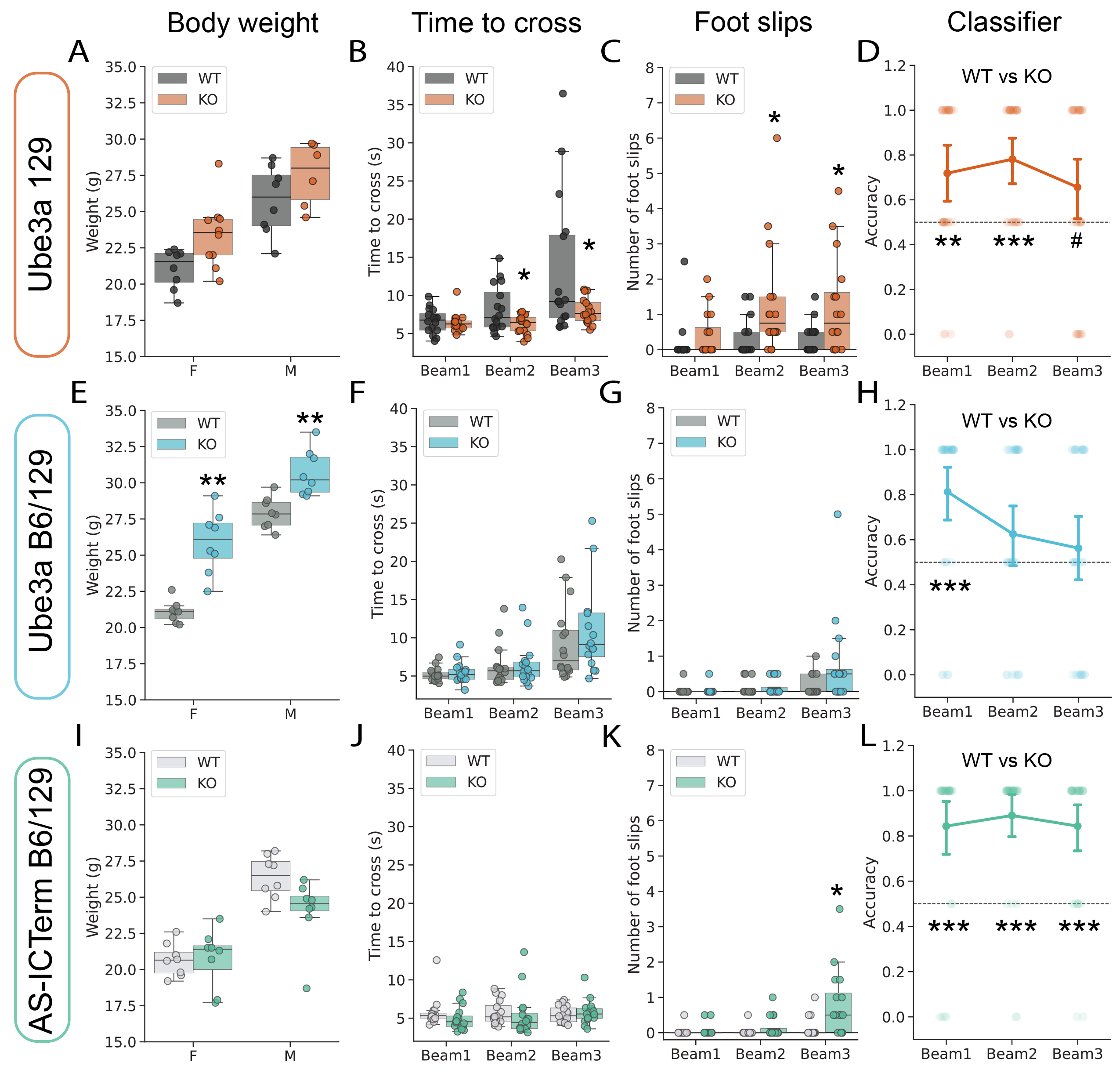
Forestwalk discriminates transgenic mouse models for Angelman syndrome in beam walking. **(A)** Weight distribution in female (F) and male (M) Ube3a 129 mice (KO) and wild-type controls (WT). Significant effect of Genotype: Two-way ANOVA, F(1,28) = 7.96, p = 0.009, and Sex: F(1, 28) = 34.62, p < 0.0001. **(B)** Time to cross the 3 different beams. Significant effect of Genotype: ANCOVA, F(1,89) = 5.32, p = 0.02, Beams: F(2,89) = 9.08, p = 0.0003 and Genotype*Beam interaction: F(2,89) = 3.1, p = 0.05. **(C)** Number of foot slips on the 3 different beams. Significant effect of Genotype: ANCOVA, F(1, 89) = 8.85, p = 0.004. **(D)** Classification accuracies of WT vs. KO mice per beam in the Ube3a 129 line. Dotted line indicates chance (0.5). Discrimination between genotypes was observed across all 3 beams. One-sample Wilcoxon test: Beam 1, W = 147, p = 0.0026; Beam 2, W = 121, p = 0.027; Beam 3: W = 125, p = 0.06. **(E)** As for A, but in Ube3a B6/129 mice. Significant effect of Genotype: Two-way ANOVA, F(1,28) = 51.56, p < 0.0001, and Sex: Two-way ANOVA, F(1, 28) = 119.82, p < 0.0001. **(F)** As for B, but in Ube3a B6/129 mice. Significant effect of Beam: ANCOVA, F(2,89) = 9.84, p = 0.0001, but no effect of Genotype, and no Beam*Genotype interaction. **(G)** As for C, but in Ube3a B6/129 mice. Significant effect of Beam: ANCOVA, F(2,89) = 5.62, p = 0.005, but no effect of Genotype, and no Beam*Genotype interaction. **(H)** As for D, but in Ube3a B6/129 mice. Discrimination between genotypes was observed in Beam 1 only. One-sample Wilcoxon test: Beam 1, W = 290, p = 0.0002. **(I)** As for A, but in AS-ICTerm mice. Significant effect of Sex: Two-way ANOVA, F(1,28) = 49.92, p < 0.0001, but no Genotype or Sex*Genotype interaction. **(J)** As for B, but in AS-ICTerm mice. No effect of Beam, Genotype, or Beam*Genotype interaction. **(K)** As for C, but in AS-ICTerm mice. Significant effect of Genotype: ANCOVA, F(1, 89) = 6.62, p = 0.012, and Beam: F(2, 89) = 9.33, p = 0.0002. **(L)** As for D, but in AS-ICTerm mice. Discrimination between genotypes was observed across all 3 beams. One-sample Wilcoxon test: Beam 1, W = 341, p < 0.0001; Beam 2, W = 375, p < 0.0001; Beam 3: W = 297, p < 0.0001. Sample size = 8 mice per sex & genotype (i.e. n=16 mice per genotype). Box plots show median ± 95% CI, point plots show mean ± 95% CI. ANCOVA was performed to adjust for body weight, with both beam and genotype as the factors of interest. Only significant comparisons with control animals are shown. # p<0.1, * p < 0.05, ** p < 0.01, *** p < 0.001.

In the beam walk test, Ube3a 129 knockout (KO) mice required less time to cross the more challenging Beams 2 and 3 (**Fig. 2B**) and performed more foot slips (**Fig. 2C**) compared to their wild-type (WT) counterparts. However, further in-depth phenotyping by Forestwalk also identified differences between KO and WT mice on the least challenging Beam 1, underscoring the increased sensitivity of our method (**Fig. 2D**). Ube3a B6/129 KO mice did not exhibit significant impairments on classical beam walk

endpoints compared to WT mice (**Fig. 2F,G**). In contrast, the RFC successfully distinguished between the two genotypes on Beam 1, again highlighting behavioral differences not captured by classical endpoints alone (**Fig. 2H**). In the case of AS-ICTerm B6/129 KO mice, the time to cross beams was comparable to WT mice (**Fig. 2J**), but mutant mice made more foot slips on the most challenging Beam 3 (**Fig. 2K**). In this strain, the RFC identified significant differences between genotypes across all three beams (**Fig. 2L**). Collectively, these findings demonstrate how the Forestwalk workflow enables the detection of differences between experimental study groups in the beam walk task, that would have gone unnoticed if using classical endpoints alone.

We next sought to understand which behavioral traits were responsible for genotype-level differences detected by Forestwalk, by taking a detailed look at body posture and prioritized features from the machine learning workflow. As significant differences between WT and KO mice were detected on Beam 1 for all strains, the subsequent analysis focused on this specific beam. For Ube3a 129 mice, no differences were seen in posture or tail position in the skeletal representations of WT and KO mice in this line (**Fig. 3A and S2 Fig A,D**). Correspondingly, the most critical features identified in Forestwalk were predominantly associated with variances in distances or angles. This suggests that the distinction between KO and WT Ube3a 129 mice lies rather in the variability of these features, but not their mean values (enrichment analysis p = 0.06, 41% enrichment). Indeed, this is exemplified by the variance of the eye y-coordinate being the most significant feature identified in Forestwalk. The time-series plot of the eye y-coordinate in KO mice displayed more pronounced and irregular fluctuations relative to WT mice; patterns often linked to compromised balance and increased incidence of missteps (**Fig. 3D** and see example video in **S5 File**).

**Fig 3.**
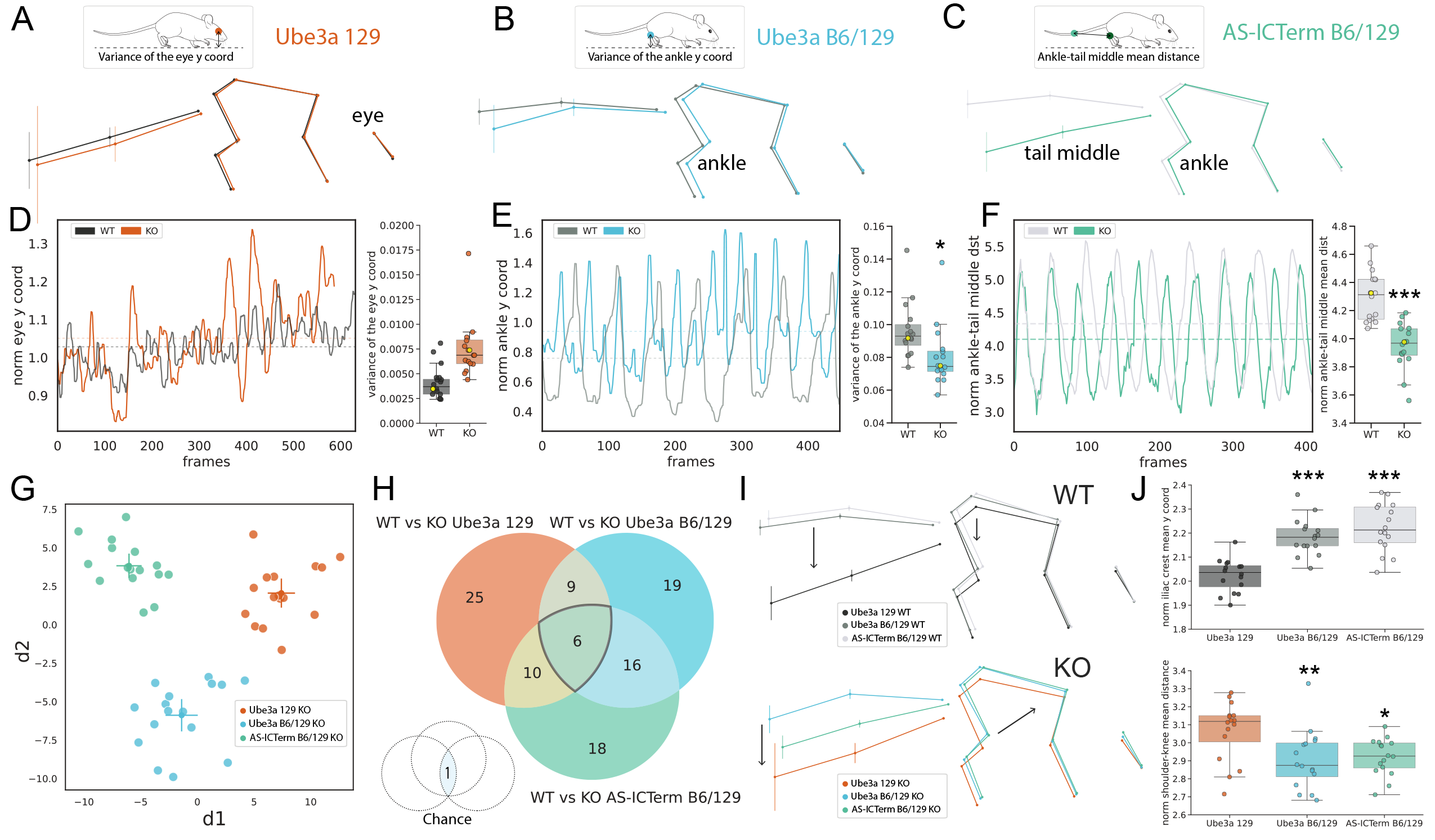
Detailed comparisons of motor coordination and balance amongst different transgenic mouse models for Angelman syndrome are enabled by Forestwalk. **(A-C)** Schematics of the average skeletal representation for each mouse line. Vertical bars show y-axis variance for each body part. The insert on top highlights the most important features discriminating KO from WT mice in each line. **(D)** Traces of normalized eye y-coordinates during a 16 mm square beam (Beam 1) walking trial are shown for representative animals of Ube3a 129 KO (orange) and WT (dark gray) genotypes. The associated box plot shows the variance of the eye y-coordinate. The data points of representative animals shown in the traces are highlighted in yellow. **(E)** As for D, but with normalized ankle y-coordinates for Ube3a B6/129 KO (blue) and WT (gray) genotypes. Significant difference between genotypes: ANCOVA, F(1,29) = 5.55, p = 0.03. **(F)** As for D, but with normalized ankle-tail middle distance for AS-ICTerm B6/129 KO (blue) and KO (gray) genotypes. Significant difference between genotypes: ANCOVA, F(1,29) = 33.85, p < 0.0001. **(G)** 2D plot reporting the first two discriminants (d1,d2) resulting from LDA on KO animals from the Ube3a 129, Ube3a B6/129 and AS-ICTerm B6/129 mouse lines. **(H)** Venn diagram illustrating feature overlap amongst the three different transgenic mouse lines. A significant commonality of prioritized features is observed vs. chance (assessed by bootstrapping with 100,000 repetitions). **(I)** Skeletal representations shown in A-C are replotted and grouped by either KO or WT genotype. **(J)** Top box plot: Elevated y-coordinate positions of the iliac crest in Ube3a B6/129 WT and AS-ICTerm B6/129 WT mice compared to Ube3a 129 WT mice. Bottom box plot: Shorter shoulder-to-knee distances in Ube3a B6/129 KO and AS-ICTerm B6/129 KO mice relative to Ube3a 129 KO. Sample size = 16 mice per genotype. Box plots show median ± 95% CI. 2D visualization of LDA shows centroid ± 95% CI. ANCOVA was performed to adjust for body weight, with genotype as the factor of interest. * p < 0.05, *** p < 0.001.

In Ube3a B6/129 mice, visual inspection of skeletal overlays suggested an alteration in body posture. Ube3a B6/129 KO mice displayed a reduced overall body length and a more compact posture compared to WT mice (**Fig. 3B**). This observation was confirmed by a significant decrease in the average distance between the shoulder and the knee in Ube3a B6/129 KO animals compared to WT littermates (**S2 Fig. B,E**). Analysis of prioritized features from Forestwalk revealed no overall category enrichment, for example in feature variance, distances or mean positions, yet the variance in the ankle y-coordinate emerged as the most significant feature (**Fig. 3B**). Time series analysis of representative subjects showed that Ube3a B6/129 KO mice had smaller, yet consistent, ankle y-coordinate fluctuations, implying a distinct and more constrained range of ankle movement relative to WT mice (**Fig. 3E** and see example video in **S5 File**).

Finally, AS-ICTerm B6/129 mice exhibited alterations in body posture akin to those observed in Ube3a B6/129 mice (**Fig. 3C**). However, in this instance, only a trend was observed toward a reduction in

the distance from the shoulder and the knee, without reaching statistical significance (**S2 Fig. C**). This implies that AS-ICTerm B6/129 KO mice may also have a tendency for a more compact posture, but the difference is less obvious than in Ube3a B6/129 mice. Notably, KO mice of this strain displayed a distinct tail posture, with the tail positioned considerably lower than WT counterparts (**Fig. 3C and S2 Fig. F**). Indeed, the most prominent feature identified in Forestwalk was the average distance between the ankle and the midpoint of the tail, which was confirmed as significantly shorter between WT and Ube3a B6/129 mutant mice (**Fig 3F**), and likely reflects the substantial change in the overall tail position (see example video in **S5 File**).

With an understanding of phenotypic differences segregating KO and WT mice for individual transgenic models, we next used LDA to further explore similarities among the three sets of KO mice. The analysis along the first discriminant axis (d1) allowed us to observe a pattern of separation among the KO mice from different strains. This pattern ranged from AS-ICTerm B6/129 KO to Ube3a B6/129 KO and then to Ube3a 129 KO, indicating that AS-ICTerm B6/129 KO mice are phenotypically closer to Ube3a B6/129 KO mice than to Ube3a 129 KO mice (**Fig. 3G**). On the other hand, the second discriminant axis (d2) clearly distinguished Ube3a B6/129 KO mice from both AS-ICTerm B6/129 KO and Ube3a 129 KO mice. This separation implies that although these strains differ in some behavioral characteristics, they also share other traits that set them apart from the Ube3a B6/129 strain, highlighting the unique behavioral signature of each mouse line (**Fig. 3G**). Consistent with this view, a comparison of prioritized features distinguishing WT from KO animals in each mouse line revealed a greater overlap between Ube3a B6/129 and AS-ICTerm B6/129 mice (22 shared features; **Fig. 3H**) than with Ube3a 129 mice (15 shared features with Ube3a B6/129 mice, 16 with AS-ICTerm B6/129 mice; **Fig. 3H**).

A comparison of prioritized features that distinguished KO from WT mice across all strains identified 6 that were common (**Fig. 3H**). Notably, 3 of these common features were related to tail position or point variance and may reflect balance disturbances common to the different mutant strains (**S2 File**). These shared features did not include postural-related measures, suggesting that postural changes may be specific to the B6/129 background, rather than the genetic perturbations common to the KO animals. Indeed, comparison of skeletal representations of all WT mice (**Fig. 3I**), revealed similar body posture between WT mice from the Ube3a B6/126 and AS-ICTerm B6/129 strains, that was distinct from WT Ube3a 129 mice. The higher body posture of the B6/129 strains vs. Ube3a 129 WT mice was confirmed by statistical comparisons of the average y-coordinate of the iliac crest (**Fig. 3J**). Meanwhile, comparison of skeletal representations for all KO mice (**Fig. 3I**), similarly revealed elevated posture in KO mice from the Ube3a B6/126 and AS-ICTerm B6/129 strains, that was distinct from KO Ube3a 129 mice. However, KO mice from the mixed B6/129 background also displayed a more compact body posture on the beam than Ube3a 129 KO mice, which was evidenced by a shorter mean shoulder-to-knee distance (**Fig. 3I,J**), and may reflect the additional impact of genetic modifiers on posture in these mice. Collectively, these findings illustrate how the Forestwalk method, utilizing intricate skeletal mappings and prioritized feature extraction, can reveal shared and strain-unique motor deficits across different transgenic models.

### Phenotypes identified by Forestwalk replicate across independent experiments

A potential concern for our approach was that prioritized features and postural phenotypes identified in one experiment may not be sufficiently generalizable or reproducible between similar experiments. To empirically test the robustness and transferability of Forestwalk, we replicated our study with a new cohort of animals, specifically selecting the AS-ICTerm B6/129 mice due to their minimal representation in prior published research.

As per Cohort 1 of AS-ICTerm B6/129 mice, no discernable differences in body weight were observed between WT and KO genotypes in the new Cohort 2 (**Fig. 4A** *cf.* **Fig 3I**). Consistent results were also seen in the classical beam walk endpoints, with no genotype difference in time to cross the beam in Cohort 2 (**Fig. 4B** *cf.* **Fig. 3J**), and an increase in the number of foot slips by AS-ICTerm B6/129 KO mice vs. WT controls only on the most challenging Beam 3 (**Fig. 4C** *cf.* **Fig 2K**). Likewise, the RFC effectively discriminated between KO and WT mice across all 3 beams (**Fig. 4D** *cf.* **Fig. 2L**). To understand if the RFC had prioritized new features to differentiate KO and WT mice in Cohort 2 than those used for Cohort 1, a comparison of prioritized features was made. Notably, prioritized features used by independent classifiers in the two experiments showed significant overlap (**Fig. 4E**). Comparing the average skeletal positions between genotypes for both cohorts revealed consistent alterations in tail positioning and animal posture (**Fig. 4F,G**). These observations were reflected by significant genotype differences in prioritized features used by the classifiers in both cohorts, including the ‘mean distance from the ankle to the tail’s midpoint’, and the ‘mean y-coordinate of the tail’s midpoint’ (**Fig. 4H,I**). Further highlighting the robustness of prioritized features to describe the genotype-specific behavior across different experiments, an RFC trained only on Cohort 1 could accurately distinguish KO from WT mice in Cohort 2 (**Fig. 4J**). Finally, LDA, using all 395 engineered features from both experiments on Beam 1, revealed that the first discriminant (d1) mostly reflected differences between genotypes, while the second discriminant reflected batch effects (d2; **Fig. 4K**). Collectively this replication experiment identifies a stable postural trait in AS-ICTerm B6/129 mice that could be the subject of further investigation, for example in the context of genetic rescue experiments. Moreover, while batch effects are present within the total feature set, Forestwalk appears to not overfit the data and can be used to identify robust, generalizable and biologically meaningful features that exhibit discriminative power between independent experiments.

**Fig 4.**
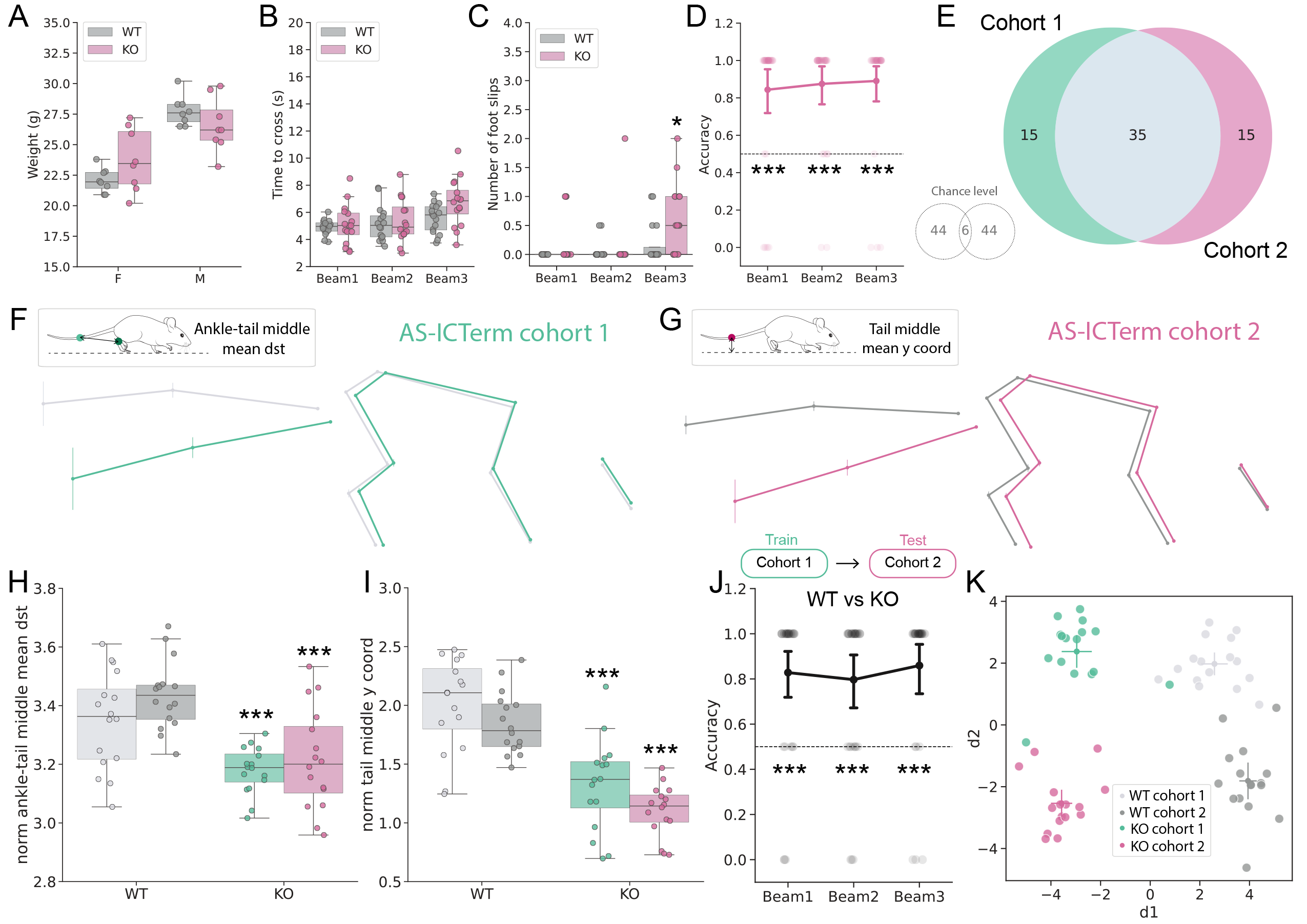
Phenotypes identified by Forestwalk are stable between independent cohorts of AS-ICTerm B6/129 mice. **(A)** Weight distribution in female (F) and male (M) AS-ICTerm B6/129 knock-out (KO) and wild-type controls (WT) from Cohort 2. Significant effect of Sex: Two-way ANOVA, F(1,28) = 41.36, p < 0.0001, and Sex*Genotype interaction: F(1,28) = 4.6, p = 0.04. **(B)** Time to cross the 3 different beams. No significant effect of Genotype. **(C)** Number of foot slips per beam. Significant effect of Genotype: ANCOVA, F(1,89) = 7.04, p = 0.009: Tukey’s test: p = 0.029 for Beam 3 in KO vs WT mice. **(D)** Classification accuracies of WT vs. KO mice in Cohort 2 per beam. A significant discrimination was achieved in all beams (One-sample Wilcoxon test, Beam 1: W = 341, p < 0.0001; Beam 2: W = 348, p < 0.0001; Beam 3: W = 400, p < 0.0001). **(E)** Venn diagram showing number of prioritized features shared by AS-ICTerm B6/129 Cohort 1 and 2. The number of features in common was significantly different from chance, as shown in the insert (p < 0.0001, assessed by bootstrapping with 100,000 repetitions). **(F)** Visualization of skeletons per genotype for AS-ICTerm B6/129 Cohort 1 and **(G)** for Cohort 2. Inserts show the most important features for the respective cohorts. **(H)** Comparison of distance between the ankle and the tail midpoint in both cohorts. Significant effect of Genotype: ANCOVA, F(1, 59) = 82.64, p < 0.0001. Tukey’s post-hoc test: Cohort 1 KO vs Cohort 1 WT p < 0.0001 (mean difference = 0.35, 95% CI: [0.17, 0.53]), Cohort 1 WT vs Cohort 2 KO p < 0.0001 (mean difference = -0.43, 95% CI: [-0.61, -0.24]), Cohort 2 WT vs. Cohort 2 KO p < 0.0001 (mean difference = -0.50, 95% CI: [-0.68, -0.32]), Cohort 2 WT vs. Cohort 1 KO p < 0.0001 (mean difference = 0.43, 95% CI: [0.25, 0.61]). **(I)** Comparison of tail middle y-coordinate in both cohorts. Significant effect of Genotype: ANCOVA, F(1, 59) = 74.91, p < 0.0001 and of Cohort: ANCOVA, F(1,59) = 7.8, p = 0.007. Tukey’s post-hoc test: Cohort 1 WT vs Cohort 1 KO p < 0.0001 (mean difference = 0.67, 95% CI: [0.37, 0.98]), Cohort 1 WT vs Cohort 2 KO p < 0.0001 (mean difference = -0.90, 95% CI: [-1.21, -0.60]), Cohort 2 WT vs Cohort 2 KO p < 0.0001 (mean difference = 0.75, 95% CI: [0.44, 1.05]), Cohort 2 WT vs Cohort 1 KO p < 0.0001 (mean difference = 0.52, 95% CI: [0.21, 0.83]). **(J)** Classifier accuracy when trained on the Cohort 1 dataset, and tested to discriminate WT vs KO mice in Cohort 2. Significant discrimination between genotypes observed in all Beams. One-sample Wilcoxon test, Beam 1: W = 294, p < 0.0001; Beam 2: W = 247, p < 0.0001; Beam 3: W = 345, p < 0.0001. **(K)** 2D plot reporting the first two discriminants (d1,d2) resulting from LDA on AS-ICTerm B6/129 Cohorts 1 and 2. Sample size = 8 mice per sex, 16 mice per genotype in each cohort. Box plots show median ± 95% CI. 2D visualization from LDA shows centroid ± 95% CI. ANCOVA was performed to adjust for body weight, with genotype as the factor of interest. Only significant comparisons with control animals are shown. * p< 0.05, *** p<0.001.

### Forestwalk identifies phenotypic differences in heterozygous and homozygous GAT1 knockout mice

In a final assessment of Forestwalk to detect genotype/phenotype differences, we turned to the GAT1 knockout mouse (B6.129S1-Slc6a1^tm1Lst^/Mmucd; (47)). This line has been used to model SLC6A1 SLC6A1-related neurodevelopmental disorder, which is characterized by epilepsy, intellectual disability, autism spectrum disorders, and motor impairments (48). Homozygous GAT1 KO mice (HO) show robust behavioral impairments, including hypoactivity, tremors, and coordination deficits (49). By contrast, heterozygous GAT1 KO mice (HE) are reported to show no overt behavioral alterations (49), despite the presence of intermittent epileptiform activity on EEG (50), and the fact that human carriers of single GAT1 mutations exhibit severe symptoms (51). Thus, here we aimed to further validate the Forestwalk approach to confirm severe motor impairments in GAT1 HO mice, while also exploring its potential to reveal yet undocumented motor deficits in GAT1 HE KO mice.

As previously reported (49), GAT1 HO mice had significantly lower body weight than either WT or HE animals (**Fig. 5A**). For classical beam walk endpoints, a significant increase in the time to cross the beam and in the number of foot slips was seen in HO animals vs. WT and HE mice across all beams (**Fig. 5B,C**). As also expected from prior literature (49), GAT1 HE KO mice showed no obvious difference in these classical endpoints as compared to WT mice (**Fig. 5B,C**). The RFC could well discriminate between HO and WT mice across all 3 beams (**Fig. 5E**), further corroborating the strong motor phenotype reported for homozygous GAT1 KO mice. Surprisingly, the RFC also achieved significant discrimination between WT and HE mice in Beam 1 (**Fig. 5D**), suggesting that motor differences are present in heterozygous GAT1 KO mice under specific test conditions, but that classical endpoints are not sensitive enough to reveal them.

**Fig 5.**
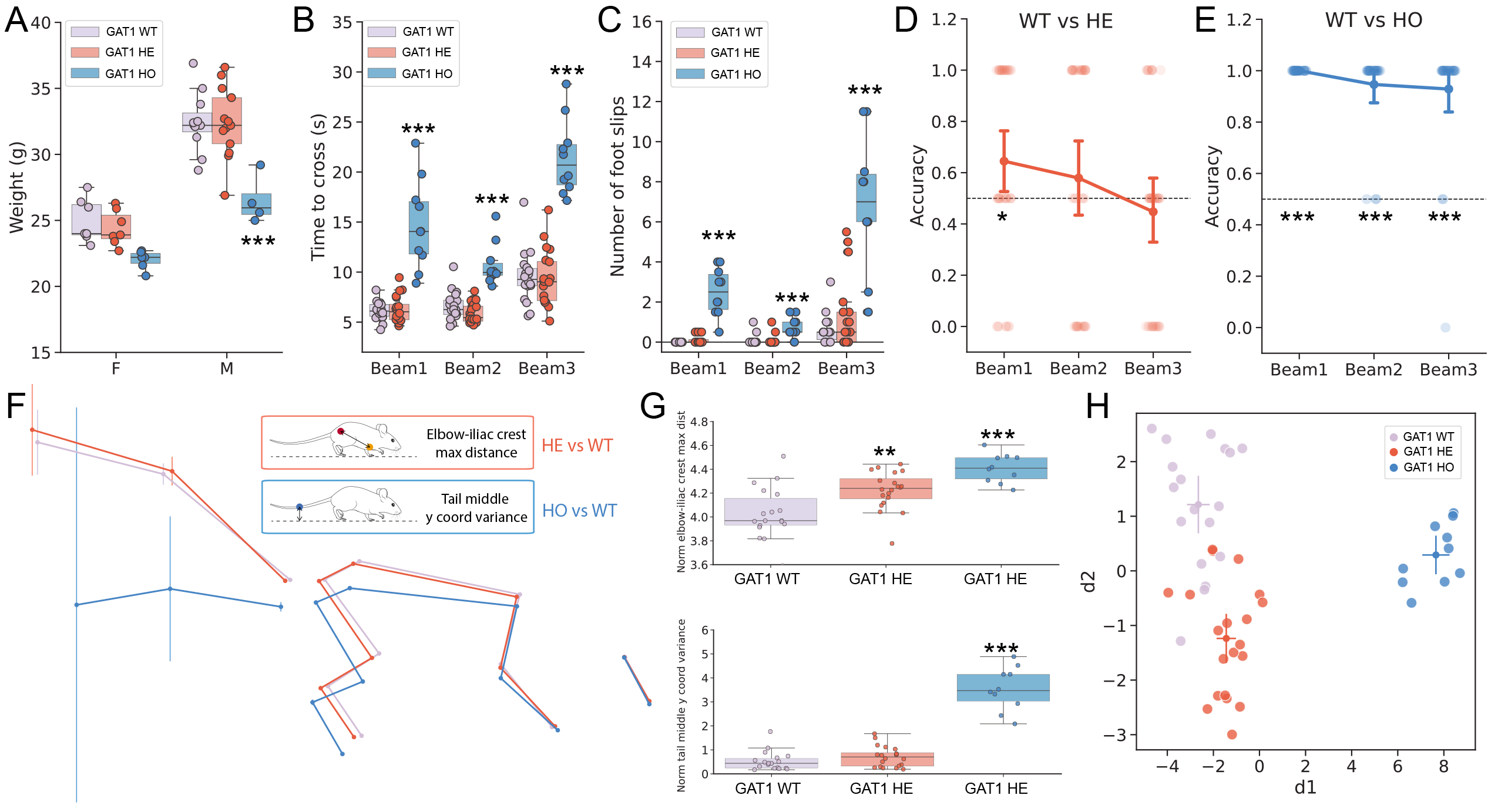
Forestwalk reveals gene-dosage effects on beam walk posture in GAT1 knock-out mice. **(A)** Weight distribution in female (F) and male (M) GAT1 knock-out heterozygous (HE) or homozygous (HO) mice, and wild-type littermate controls (WT). Significant effect of Genotype: Two-way ANOVA, F(2,42) = 15.39, p < 0.0001. Tukey’s test: male HO vs. male WT p = 0.0002, male HE vs male WT p = 0.0002. **(B)** Time to cross the beam. Significant effect of Genotype: ANCOVA, F(2,134) = 126.92, p < 0.0001), and Beam: ANCOVA, F(2,134) = 60.20, p < 0.0001, and Genotype*Beam interaction: ANCOVA, F(4, 134) = 11, p < 0.0001. Tukey’s test: GAT1 HO vs. WT mice on Beam 1: p < 0.0001; Beam 2: p < 0.0001 and Beam 3: p < 0.0001, or GAT1 HO vs. HE mice on Beam 1: p < 0.0001; Beam 2: p < 0.0001 and Beam 3: p < 0.0001. **(C)** Number of foot slips. Significant effect of Genotype: ANCOVA, F(2, 134) = 62.15, p < 0.0001, of Beam: ANCOVA, F(2, 134) = 40.36, p < 0.0001, and Genotype*Beam interaction: ANCOVA, F(4, 134) = 22.32, p < 0.0001. Tukey’s test: GAT1 HO vs WT mice on Beam 1: p < 0.0001; Beam 2: p = 0.0094 and Beam 3: p < 0.0001, or GAT1 HO vs or HE mice on Beam 1: p < 0.0001; Beam 2: p = 0.0068 and Beam 3: p < 0.0001. **(D)** Classifier accuracy in discriminating GAT1 HE from WT mice. Significant discrimination achieved in Beam 1 (One-sample Wilcoxon test: W = 143, p = 0.043). **(E)** Classifier accuracy in discriminating GAT1 HO from WT mice. Significant discrimination achieved in all beams (One-sample Wilcoxon test, Beam 1: W = 406, p < 0.0001; Beam 2, W = 325, p < 0.0001; Beam 3, W = 324, p < 0.0001). **(F)** Visualizations of skeletons per genotype. Inserts show the most important features in discriminating between WT and HE mice, or WT and HO mice. **(G)** Top box plot: Maximum distance between the elbow and the iliac crest per genotype. Significant effect of Genotype: ANCOVA, F(2, 44) = 14.03, p < 0.0001. Tukey’s post-hoc test: HE vs WT p = 0.004 (mean difference = -0.19, 95% CI: [-0.33, -0.05]), HO vs WT p < 0.0001 (mean difference = -0.38, 95% CI: [-0.55, -0.21]), HO vs HE p = 0.02 (mean difference = 0.19, 95% CI: [0.02, 0.35]). Bottom box plot: Variance of the y-coordinate of the tail midpoint per genotype. Significant effect of Genotype: ANCOVA, F(2, 44) = 76.52, p < 0.0001. Tukey’s post-hoc test: HO vs WT p < 0.0001 (mean difference = -3.01, 95% CI: [-3.53, -2.48]), HO vs HE p < 0.0001 (mean difference = 2.83, 95% CI: [2.31, 3.34]. **(H)** 2D plot reporting the first two discriminants (d1,d2) resulting from LDA on GAT1 mice. Sample size = 18 WT (7 females, 11 males), 20 (7 females, 13 males) HE and 10 HO (6 females, 4 males). Box plots show median ± 95% CI. 2D visualization from LDA shows centroid ± 95% CI. ANCOVA was performed to adjust for body weight, with genotype as the factor of interest. Only significant comparisons with control animals are shown. ** p< 0.01, *** p<0.001.

We explored postural differences in GAT1 mutant mice in more detail and focused on Beam 1 where both HO and HE mice were discriminated from WT controls by the RFCs. The averaged skeletal alignments and prioritized features revealed substantial changes in GAT1 HO mice vs WT controls. In particular, HO mice assumed an extended posture (**Fig. 5F)**, evidenced by an increased elbow-to-iliac crest distance (**Fig. 5G and S3 Fig. A**). HO mice also displayed a lower center of gravity, with a reduced iliac crest height (**S3 Fig. B**) and lower hindpaw position on the beam’s edge (**S3 Fig. C**), likely to achieve increased stability while traversing the beam (**Fig. 5F** and see example video in **S7 File**). These postural changes were accompanied by extensive movements in the tail of GAT1 HO mice. Indeed, variance in the tail middle point was a top-ranked prioritized feature distinguishing WT and HO mice (**Fig. 5G**). Similarly, GAT1 HE mice also showed a modest extension in body length when compared to WT mice (**Fig. 5F, G and S3 Fig. A**), but this did not coincide with a significant change in the center of mass (**S3 Fig. B**). The observed elongation in posture was reflected in the prioritized features, such as the maximum distance between the iliac crest and the elbow (**Fig. 5G**). Finally, LDA visualization further substantiated differences amongst genotypes. The first discriminant (d1) effectively separated HO mice, while the second discriminant (d2) distinguished HE mice from WT mice, confirming the presence of a balance phenotype in these animals (**Fig. 5H and S3 Fig. D**). Collectively, these findings underscore the ability of Forestwalk to identify genotype/phenotype relationships in motor function and opens the door for further study of brain-behavior relationships in such genetic models of human disease.

## Discussion

Investigating motor function through animal models is essential to understand brain function in health and disease and to develop potential therapeutic strategies (1). In this context, behavioral tests such as the beam walk provide an invaluable, controlled and quantifiable way to evaluate coordination and balance in rodents, and also humans (4,21,23). However, classical endpoints used in the rodent beam walk, including the number of foot slips and time taken to cross the beam, do not fully capture the repertoire of behaviors observed during the task. Additionally, manual scoring of classical endpoints gives rise to high variability between different observers, which limits the reproducibility of findings. Here, using machine learning-based data capture and analytical tools, we have revisited the rodent beam walk to ask what additional information can be obtained from this ethologically relevant test. We developed a new analysis workflow for the beam walk, named Forestwalk, and were able to identify previously hidden effects of pharmacological treatment and unreported phenotypes in different mouse models used to research neurodevelopmental disorders. Taken together, this new approach can advance research into brain mechanisms controlling posture and balance.

An initial objective of our study was to automate the detection of classical endpoints used in the beam walk, namely the number of foot slips and time to cross the beam. This was achieved with Forestwalk, and thus ensures a standardized and reproducible scoring approach, as compared to human-raters who are prone to inter- and intra-rater variability. Forestwalk also results in a considerable time saving in comparison to manual scoring. Automated scoring of foot slips and time to cross of > 180 videos from one experiment, with a total recording duration of ∼66 minutes, took only ∼4 minutes when using a high-performance computing cluster. In contrast, human-raters typically score videos at a reduced playback speed or even frame-by-frame, so would require considerably more time to score the same data set. Running the entire Forestwalk workflow on the same experiment took ∼65 minutes and, given the high number of features explored in each video, manual annotation and analysis of such an extensive feature set is unlikely to be even attempted by a human-rater.

A further objective of our study was to improve the sensitivity of the beam walk to reveal differences between experimental groups. In one example using diazepam, we identified treatment effects with a low dose of diazepam in the expanded feature set of Forestwalk that were not seen when using classical endpoints alone. In pharmaceutical drug discovery, obtaining such insights are critical. Accurate dose-response relationships are necessary to prioritize compounds for further development, to establish the therapeutic window between efficacy and safety, and to inform dose selection for future clinical studies. Moreover, studies using beam walking typically claim successful rescue of model phenotypes when foot slips are monitored and recovered to control levels or claim absence of a motor phenotype or treatment effect when no difference in foot slips occurs. Our study clearly demonstrates that subtle but relevant changes in posture and balance can still occur even in the absence of changes in foot slips. Thus, caution should be taken when using classical endpoints alone in drawing conclusions from the beam walk test.

A final objective for our research was to address the limited specificity of the beam walk task to discriminate fundamentally different experimental conditions. We focused on comparing transgenic mouse strains used to study Angelman syndrome (AS), which is a genetic disorder affecting the nervous system, and is characterized by developmental delay, intellectual disability, profound speech impairment, and problems with movement and balance (39,52). The rotarod is most often used to study balance and coordination in mouse models for AS, and has identified impaired performance across several transgenic lines (42,43,53) and been used to assess therapeutic interventions (54–58). A particular strength of the beam walk in comparison to the rotarod is that it enables an assessment of *how* the animal performs the test, not only *if* the animal performs it (9). Such qualitative differences in performance, which can now be reliably quantified in the Forestwalk workflow, are likely to facilitate a more precise understanding of mechanisms linked to balance control in AS and would provide a more granular view on therapeutic efficacy. Indeed, we were able to identify common and unique postural traits associated with each transgenic line under investigation, which may speak to the genetic commonalities (e.g., loss of ube3a) and differences (e.g., background strains) amongst the lines. Similarities were identified between Ube3a B6/129 and AS-ICTerm mice, which may reflect their shared genetic background. Across all three transgenic lines, 6 shared features were identified, of which 3 were related to tail position or movement, consistent with balance disturbances. Moreover, prioritized features distinguishing transgenic mice from controls were stable between independent cohorts of AS-ICTerm B6/129 mice, suggesting that they could serve as defined endophenotypes of the line. In light of our findings, we suggest that further attention should be given to the beam walk in the characterization of posture and balance disturbances in rodent lines used for AS research and for future therapeutic testing.

A concern for machine learning models such as those used in our study is that they may have limited generalizability beyond the dataset and conditions on which they were initially developed. Although our DLC model was trained to detect key points on mice of different ages, weights and fur color, other variables such as room lighting conditions, the camera used, camera positioning, and the beam equipment itself, could impact on the accuracy and transferability of the model (29). To overcome this, establishing a new DLC key-point model, or re-training the model used in our study with examples provided from a new experimental context is highly recommended. We further explored the generalizability of our approach by performing a replication experiment with AS-ICTerm B6/129 mice and found that a classifier trained on one cohort of animals could well-predict the genotype of mice in a second fully independent cohort. Thus, this cross-cohort validation suggests that our model has not been overfitted to the initial group and has indeed captured essential features that differentiate the genotypes. Interestingly, when looking at the entire feature set from these experiments, we found that its information content not only pertained to the primary experimental factors under investigation (i.e., wild type vs. transgenic mouse differences), but also identified global batch differences (i.e., cohort 1 vs. cohort 2). While potential sources of variability (e.g., beam type, time of day, room lighting conditions) were controlled between cohorts, it is clear that other factors that may contribute to between-batch variability are less well understood (e.g., slow oscillatory activity in laboratory housed mice; (59)). We suggest that the Forestwalk approach, given its improved sensitivity, can also be used to further study the influence of such variables on experimental outcomes.

To further understand if the sensitivity of Forestwalk was appropriately calibrated, we conducted studies in GAT1 knockout mice (47), a useful model for the study of SLC6A1-related neurodevelopmental disorder (48,51). Homozygous GAT1 KO mice exhibit a range of motor disorders, including gait abnormalities, constant tremor, reduced rotarod performance and reduced locomotor activity in the homecage (49). By contrast, extensive study of heterozygous GAT1 KO mice has failed to reveal pronounced behavioral phenotypes (49), despite the fact that heterozygous mice have only intermediate GABA uptake capacity (49) and exhibit frequent spike-wave-discharges measured by electrocorticography (50). Consistent with published literature, we identified pronounced differences in GAT1 HO vs. WT controls with both classical endpoints and using the expanded feature set of Forestwalk. For GAT1 HE mice, classical endpoints did not differ from WT controls. Nevertheless, the classifier was able to successfully categorize GAT1 HE mice in the first beam. The absence of detectable differences in GAT1 HE mice on other beams is an interesting result that requires further study but could implicate a learning effect that only manifested during exposure to the first beam. Taken together, studies with GAT1 KO mice suggest that Forestwalk generates results that are consistent with other published reports but can yet reveal subtle group differences that may have escaped detection in other measures of motor function and warrant further exploration.

Optimizing beam walk data capture and analysis has also been the focus of other groups (13,14,18,24,37). For example, Ito et al., developed a ‘simple scoring system’ with binary judgements of values, such as retention, moving forward and reaching the goal to rate beam walking performance (13). A low score thus reflected poor performance, and the approach was validated using a model of SCI with comparison to another commonly used scale in the field, namely the Basso Mouse Scale for locomotion

(60). Other groups have adopted a single-frame motion analysis (SFMA) approach in both mice (35,36) and rats (14) to assess gait deficits after spinal cord or nerve injury. In SFMA, objective parameters are obtained from individual video frames, such as the foot-stepping angle, and these parameters were found to correlate with underlying lesion volume and to track lesion severity (14). While the simple scoring method and SFMA are both straightforward to implement, require only cheap equipment and no specific knowledge or training of personnel (13,14), they do rely on human-raters and require significant time to score or manually annotate videos.

Like in our study, other groups have turned to DLC-based tracking to overcome the limitations of human-rater scoring of beam walking performance (18,24,37). Lang et al., established a parameter to assess rhythmicity of gait, and reported changes in beam walking performance in a mouse model for spinocerebellar ataxia that were not observed in less complex studies of natural gait (24). Meanwhile, Wan et al., deployed FluoRender architecture to extract a standard walk cycle from 2D and 3D pose-estimation, and demonstrated feasibility of the method in two mice (37). Most recently, Bidgood et al., combined DLC tracking with an automated behavioral classification tool, called Simple Behavioral Analysis (28) and identified differences in global walking dynamics in a mouse model used for Parkinson’s disease research (18). While each of these methods has its own advantages, to the best of our knowledge, Forestwalk is the only approach thus far to automate detection of foot slips and is scalable to full experimental studies. Moreover, DLC labeled key-points can be used in Forestwalk to reconstruct faithful skeletal representations of mice and combine engineered features with machine learning tools. Collectively, this workflow permits automated and detailed exploration of determinant postural differences in mice traversing the beam that has not, to the best of our knowledge, previously been accomplished.

The beam walk task is of particular interest given its direct translational application to human studies. Balance is important for many aspects of daily life of humans, including walking (61), standing from a chair (62) or climbing stairs (63). Impaired balance and gait, potentially leading to falls, is a major determinant of poor quality of life, immobilization and reduced life expectancy in older adults, and in several neurological conditions (64,65). As in our report, studies of balance in humans using beam walking are also exploring the potential for marker-less-based motion capture to overcome issues associated with maker-based systems (21). As marker-less pose estimation becomes more commonplace, it will be of great value to align and potentially harmonize analytical methods used in non-clinical and clinical settings where possible to further aid translational research.

### Limitations

There are several limitations to our study. First, we chose a single side-view camera, which precluded monitoring of both hind paws for foot slip detection. Our method could be improved by the use of an additional camera, or by placement of a mirror to obtain additional views of the mouse, as recently described in a study of stroke recovery using the ladder rung test (66). Second, human studies with marker-less based motion capture have placed emphasis on comparisons to marker-based methods for validation and comparison of accuracy of these approaches (21). We did not benchmark against marker-based methods, which place an additional welfare burden in mice. Thus, it is possible that potential bias was introduced into our analysis workflow from key-point tracking data that is difficult to objectively quantify. Third, our approach focused on mice, but the beam walk is also commonly used with rats (5,8,14). It would be of value in the future to extend the Forestwalk workflow to this species. We expect that a new key-point model would need to be trained specifically for rats, and automated foot-slip detection would need to be re-established using different threshold parameters. However, general principles of the Forestwalk workflow would also be applicable to data obtained from studies in rats. Fourth, our approach does require computational resources and programming expertise that may not be available to all laboratories. Notwithstanding, we do believe that the additional insights gained from our approach far outweigh the investment necessary to obtain these insights, both from a scientific and ethical standpoint. Fifth, we recognize that automated analysis can contribute to reducing variability and increase experimental reproducibility. However, additional factors still need to be considered (such as the equipment type, batch effects and learning effects due to procedural differences between laboratories) to ensure reliable and reproducible results from the beam walk that are beyond the scope of our work. Finally, here we focused only on the beam walk, and were unable to make direct comparison between performance in this task and other measures of balance and coordination, such as rotarod performance. Eventually, all tests for motor behavior have advantages and limitations, and it is typically recommended that a battery of measures appropriate for a particular study are deployed (14,23).

## Conclusion

Here we established a new analysis workflow for beam walking in mice, named Forestwalk, that automates the detection of classical endpoints used in the task, and delivers more sensitive and specific insights into postural control and balance. As demonstrated by our experiments with diazepam and different transgenic strains used in the study of neurodevelopmental disorders, Forestwalk opens the door to a better understanding of how the brain controls movement in health and disease.

## Materials and methods

### Animals

For pharmacology studies with diazepam, male C57BL/6J mice (Jax ID:000664, aged 14 weeks old) were obtained from Charles River Laboratories (Saint Germain sur l’Arbresle, France). Experiments in genetic models for neurodevelopmental disorders included male and female, mutant and wild-type littermates of Ube3a 129 mice (129-Ube3a^tm1Alb^/J, Jax ID: 004477; aged 8 weeks); Ube3a B6/129 hybrid mice (B6.129S7-Ube3a^tm1Alb^/J, Jax ID: 016590; aged 9 weeks); AS-ICTerm mice (C57BL/6J-Rr70^em1Rsnk^/Mmmh, 065423-MU; aged 7-8 weeks) and GAT1 mice (B6.129S1-Slc6a1^tm1Lst^/Mmucd; 000426-UCD; aged 18 weeks). All transgenic mice were bred externally (Taconic, Denmark or Charles River, Germany) and shipped to the test facility (Roche Innovation Center, Basel) after weaning. Mice were acclimated to the test facility for at least one week before starting experiments. All animals were experimentally naive prior to starting beam walk experiments.

Mice were housed in groups of 2-3 per cage (GM500, Tecniplast), unless instances of aggression between littermates required single housing. Woodchip bedding (SAFE FS 14, J. Rettenmaier & Söhne GmbH) was used in cages, with nesting material and additional in-cage enrichment items provided, which were replaced during each cage change. Mice had *ad libitum* access to food (Standard Diet; Kliba Nafag) and water in the home cage. Temperature (22°C ± 2°C) and humidity (50 ± 10 %) was controlled in housing and experimental rooms, and mice were maintained under a 12h:12h Light:Dark cycle (lights transitioning to on at 06:00), with beam walk tests conducted during the light phase.

All animal experiments were conducted in strict adherence to the Swiss federal ordinance on animal protection and welfare, as well as according to the rules of the Association for Assessment and Accreditation of Laboratory Animal Care International (AAALACi), and with the explicit approval of the local veterinary authorities.

### Beam walk test

The beam walk test was employed to evaluate rodent coordination, posture and balance (1,67). This assessment measures the ability of the rodents to traverse a series of 1 m long narrow beams of varying dimensions and shapes (16 mm square, 16 mm round, 9 mm square) to reach a goal area (a dark box containing a home cage with clean bedding material). The beams were suspended 30 cm above a bench, with a soft pad placed underneath to cushion any falls. Prior to the beam walk test, mice underwent a minimum of three training trials to ensure their ability to cross the beam. These trials progressed from first placing the mouse in close proximity to the box, to the middle of the beam, and finally at the start of the beam. Before the start of the test, mice were acclimated for 2 minutes in the goal area. In a standard experiment, mice were required to traverse the 16 mm square rod twice, followed by the 16 mm round rod twice, and finally the 9 mm square rod twice. A rest period of 15 minutes was provided between each rod. In pharmacological experiments, the protocol was slightly modified. Mice traversed each beam once, rested for a minimum of 15 minutes, and then repeated the sequence. This configuration facilitated a more rapid assessment of a specific compound’s effect. In instances where a mouse fell from the beam, a second attempt was permitted. However, a second fall resulted in the termination of all subsequent beam walk testing for that mouse on that day.

### Pharmacological treatment

On the test day, mice (n=2 per group) were randomly assigned to receive either diazepam (synthesized by F. Hoffmann-La Roche) 0.3, 1, 3 mg/kg or its vehicle (NaCl 0.9% +Tween 80 0.3%) administered intraperitoneally 30 minutes before being placed on the beam. The experiment was repeated once per week until all animals received all treatment conditions, following a within-subjects randomized cross-over design.

### Pose estimation and tracking

Behavioral videos were acquired using a Basler C-Mount acA1920-150um camera (2.3 MP resolution, 120 fps; Basler, Germany) and a HF25XA-5M. F1.6/25mm lens (Chromos Group AG, Switzerland). DeepLabCut 2.2.3 was used to track 18 key points, including 5 reference points on the beam and 13 mouse body parts including the nose, the eye, the forepaw, the elbow, the shoulder, the hindpaw, the ankle, the knee, the hip, the iliac crest, the base of the tail, the tail center and the tail tip. The final network was trained by using a total of 520 frames extracted from 26 videos for 1,030,000 iterations.

### Forestwalk workflow

The coordinates extracted by DeepLabCut were imported in Python and the points related to the beam were used to perform the automatic detection of the beam type (16 mm square beam: a small dot on the right side; 16 mm round beam: no dot; 9 mm square beam: small dot on the left side). The central 80 cm region of the beam was detected using the key points tracking the two vertical black lines displayed at the two extremities of each beam. The final error of the network was 3.11 pixels on the training set and 3.82 pixels on the test set, with the training set representing 95% of the data.

### Automated detection of time to cross and number of foot slips

The method developed for automated detection of time to cross and number of foot slips is given in the results section. Foot slip detection accuracy was evaluated by comparing an automated threshold-based method with reference annotations made by a single human scorer.

A true positive (TP) was identified when the automated method’s detection aligned with a foot slip noted by the scorer within a predetermined frame range. A false negative (FN) occurred when the scorer marked a slip that the automated method did not detect, while a false positive (FP) was a detection by the automated method not confirmed by the scorer. Various thresholds were tested to fine-tune the automated method. Performance metrics—recall (TP / (TP + FN)), precision (TP / (TP + FP)), and the F1 score—guided the selection of the optimal threshold, which was determined by the highest combined recall and precision to ensure a balanced detection of slips with minimal false identifications.

Agreement between different human scores and the automated method for foot slip detection was calculated using Cohen’s kappa. Cohen’s kappa coefficient was calculated using the ‘cohen_kappa_score’ function from Sklearn.metrics. Agreement was assessed on a per-video basis by comparing the total foot slip counts identified by each scorer for each video, providing a video-level measure of scorer consistency.

### Feature engineering

Pose estimation data collected from each behavioral video were utilized to generate a set of behavioral features. Features were obtained by calculating all the pairwise distances between the animal’s body parts throughout the video, the angles at the elbow, ankle, knee, and hip levels, and the distance of each body part from the beam surface. To account for differences in animal body size, distances were normalized by a reference, which was chosen to be the distance between the elbow and the shoulder, due to its strong correlation with body weight. To condense the information contained within these features without working with time series, we calculated the mean value, the minimum value, the maximum value, and the variance for each feature. This resulted in 395 features per video (See **S1 File 1)**, that also included the animal sex, weight, and classical endpoints of number of foot slips and time to cross.

### Random forest classification and feature selection

A Random Forest Classifier (RFC) was utilized to explore the potential of behavioral features derived from each experiment in discriminating specific biological conditions, such as variations in genotypes or doses of a particular compound. The choice of a RFC was driven by its ability to rank features based on their importance, its robustness to overfitting due to the independent training of numerous trees, and its suitability for small input datasets, which was the case in our study.

### Partitioning data into training, validation, and test sets

For each experiment, the feature datasets were initially divided into training, validation, and test sets using a leave-one-out cross-validation (LOO-CV) approach. Subsequently, Z-score normalization, derived from the training set, was applied to the training set itself as well as to the validation and test sets. This approach is advantageous as it optimizes data usage and provides a robust estimate of model performance. Given that multiple data points were collected from each subject in this study, we used a LOO-CV to evaluate the model’s performance and to prevent data leakage from the training to the validation process. During LOO-CV, all data points from a specific subject were reserved for testing, all points from a different animal were set aside for validation, and the data from all remaining subjects were used for model training. This process was iterated for all subjects within each group. As outlined in the Beam Walk Test section, each animal was permitted to attempt crossing each beam a maximum of two times. In the context of the model, each beam crossing by the same animal was considered a unique data point. For every such data point, a corresponding set of features was compiled, and an accuracy value was determined.

### Feature selection and classification

Using the training and validation data, we (i) selected the 50 most critical features to reduce multicollinearity, increase the signal-to-noise ratio, and enhance the model’s performance. Specifically, we used the Recursive Feature Elimination (RFE) method from the sklearn.feature_selection module for feature selection, with rfe_step = 0.05, and the RFC from the sklearn.ensemble module as the estimator, with default parameters except for class_weight = ‘balanced’, n_features_to_select = 50; (ii) tuned the model hyperparameters using these 50 critical features through GridSearchCV from the sklearn.model_selection module; (iii) performed a second round of feature selection using the tuned hyperparameters, and ranked all the features based on their importance, as measured by the mean decrease in impurity. Then, for every animal, we obtained a final set of prioritized features by comparing the ranked feature lists from the train/validation leave-one-out split with the same test set. We created a consensus ranked list considering both the occurrence and the rank of each feature within all lists and selected the first 50. The final hyperparameters were instead obtained by taking the mode of each hyperparameter selected in all the train/validation splits. For each test set, then, we trained a random forest classifier using the specific prioritized features and hyperparameters selected from the training step and we obtained an accuracy score ranging from 0 to 1, due to the presence of maximum two data points per animal on a specific test set.

### Linear discriminant analysis

Linear discriminant analysis (LDA) was employed as a technique to visualize the separation between different biological groups within our datasets. For our analysis, we utilized the LDA implementation provided by the scikit-learn library in Python (68), adhering to the default parameter settings. This included using the singular value decomposition approach without assigning any priors on the class proportions. All available features in the dataset were included in the LDA. The resulting two-dimensional projection was then used to assess and illustrate the degree of group distinction based on the given behavioral features. To further quantify the separability between the groups identified in our dataset, we measured the Euclidean distance from the centroid of a reference group to each sample belonging to that group to assess the intra-group variability. Then, we compared these intra-group distances with the distances between the samples belonging to a different group and the centroid of the reference group. A larger average distance between groups, as compared to the average intra-group distances, would indicate a clear separation between the two groups.

### Skeleton visualization

In order to visualize the average skeleton for each animal, we first standardized the x-coordinate of the nose to zero for all animals. We then calculated the x-coordinate for each remaining body part by using the mean distance of that body part from the nose normalized on the reference dimension. This allowed us to maintain the relative positioning of each body part while standardizing the overall skeleton size and orientation across animals. For the y-coordinate, we used the corresponding mean y-coordinate for each body part. This mean y-coordinate was also used as a feature in our analysis. To represent the variance around each body part, we added bars to each body part in the visualization. These bars only represent variance in the y-axis, as this was the primary axis of interest in our study. This method allowed us to create a clear, standardized visualization of the average skeleton for each animal, highlighting the key features and variances of interest.

### Statistical analysis & study design factors

For pharmacological studies with diazepam, beam walk performance following vehicle administration served as the control. For studies in transgenic mice, wild-type littermates served as the control group. We selected a sample size based on that typically used in the literature (i.e. minimally n=8 per group to provide sufficient power to detect differences of 10-15% in group means (4)). All mice ordered were included in the study and subsequent analysis. Mice were only excluded in case of a technical error (e.g. video recording failure). We set an exclusion criteria of 1 minute for mice to cross the beam, although in our experience no mouse exceeded this threshold. Blinding was not performed in the study - i.e. experimenters were aware of the treatment/vehicle allocation in the diazepam experiment, and mouse genotype in studies with genetic models. Differences in beam-crossing time and foot slips were analyzed using ANCOVA, with weight as a covariate for the diazepam experiment and weight and beam as covariates for other experiments in transgenic mice. Classifier accuracy against chance level (0.5) was assessed with a one-sample Wilcoxon test. Two-way ANOVA was applied to body weight data, considering genotype and sex as factors. Feature overlap significance was determined through bootstrap sampling (100,000 repetitions), and feature category enrichment was evaluated using the hypergeometric test.

## Data Access

DLC annotated key-point data from >1200 video files recorded for this publication will be made available at https://doi.org/10.5281/zenodo.11068060 at the time of publication. Data files are provided with a Minimal Metadata Set (69) to enable their future reuse and repurposing. Open- source code for Foreswalk will be made available for download at https://github.com/Roche/neuro-ForestWalk at the time of publication.

## Author contributions

F.T.: Methodology, Software, Validation, Formal Analysis, Investigation, Data Curation, Writing - Original draft, Writing - Review & Editing, Visualization.

Y-P. Z.: Conceptualization, Software, Data Curation, Writing - Review & Editing, Supervision R.N.: Conceptualization, Writing - Review & Editing, Supervision

D.R.: Conceptualization, Software, Formal Analysis, Data Curation, Writing - Review & Editing, Supervision

E.C.O’C.: Conceptualization, Writing - Original draft, Writing - Review & Editing, Methodology, Visualization, Supervision, Project administration, Internal funding & resource acquisition.

## Disclosure

All authors were employees of F. Hoffmann-La Roche AG Switzerland at the time of study conduct and original manuscript submission.

## Acknowledgements

We thank Brigitte Algeyer, Marie Haman, and Roger Wyler for technical support in data collection, and the Animal Care and Welfare Staff of IVR98 at the Roche Innovation Center, F. Hoffmann-La Roche, Basel. B6.129S1-Slc6a1^tm1Lst^/Mmucd mice were obtained under a Material Transfer Agreement from the California Institute of Technology. C57BL/6J-Rr70^em1Rsnk^/Mmmh mice were obtained under a Material Transfer Agreement from the University of Florida. We thank scidraw.io for the illustrations (https://doi.org/10.5281/zenodo.3925915).

