## Supporting information for "Forestwalk: A machine learning workflow brings new insights into posture and balance in rodent beam walking"

**Short title:**

Forestwalk: New insights from rodent beam walking

### Supporting information

### S1 Fig.

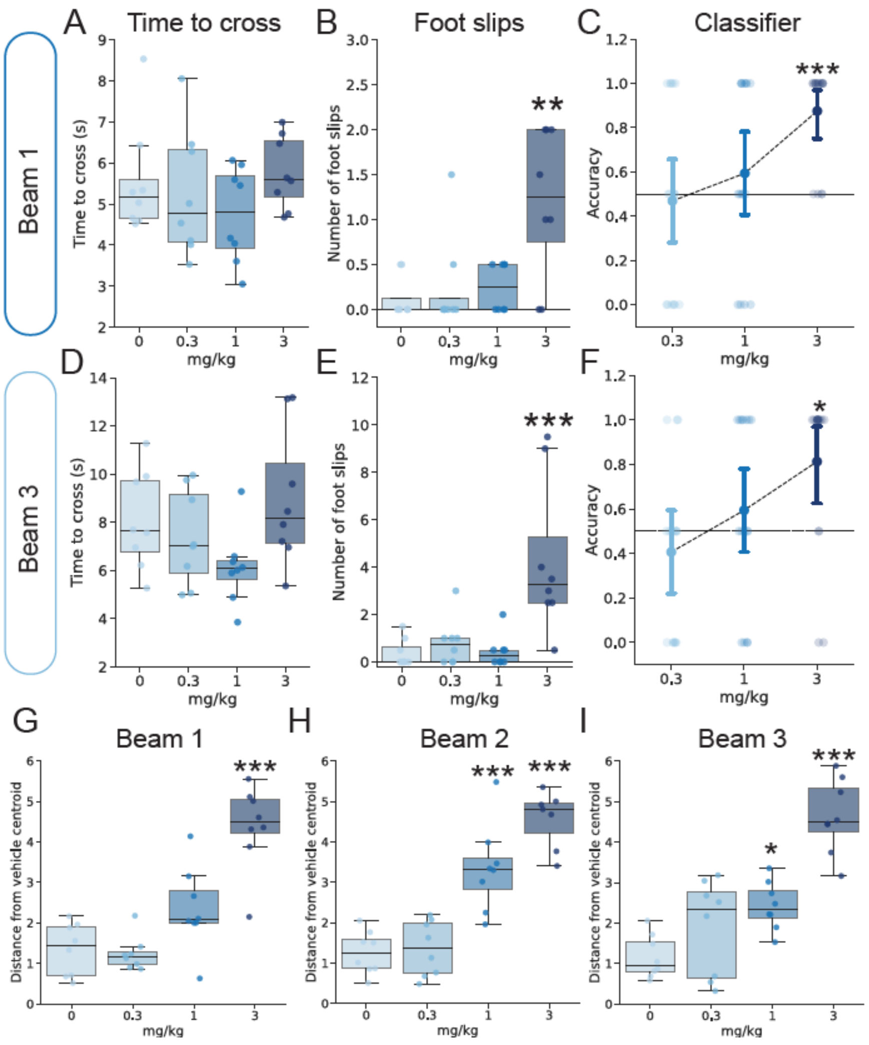


**S1 Fig. Analysis of Beam 1 and 3 data, and Euclidean distance from LDA from all beams in the diazepam experiment. (A)** Time to cross the beam (Beam 1) in the diazepam experiment. Effect of treatment: ANCOVA, $F(3, 27) = 1.13$, $p = 0.35$. **(B)** Number of foot slips (Beam 1) in the diazepam experiment. Effect of treatment: ANCOVA, $F(3, 27) = 7.38$, p(dose) = 0.0009; Tukey's post-hoc test: 3 mg/kg vs 0 mg/kg $p = 0.002$, mean difference = -1.06, 95% CI: [-1.79, -0.34]. **(C)** Classification accuracies resulting from the comparison between animals treated with diazepam and vehicle in Beam 1. Dotted line indicates chance (0.5). One-sample Wilcoxon test: 3 mg/kg vs. chance $p = 0.0005$, $W = 78$. **(D)** As for panel A, but with Beam 3. Effect of treatment: ANCOVA, $F(3, 27) = 2.60$, $p = 0.07$. **(E)** As for panel B, but for Beam 3. Effect of treatment: ANCOVA, $F(3, 27) = 10.00$, p(dose) = 0.0001; Tukey's post-hoc test: 3 mg/kg vs 0 mg/kg p = 0.0006, mean difference = -3.94, 95% CI: [-6.32, -1.56]. **(F)** As for panel C, but for Beam 3. One-sample Wilcoxon test: 3 mg/kg vs. chance p = 0.0129, W = 75. **(G)** Euclidean distance from the centroid of the vehicle cluster identified in the space of the first two discriminants following LDA analysis on Beam 1. ANCOVA, F(3, 27) = 23.96, p(dose) < 0.0001; Tukey's post-hoc test: 3 mg/kg vs 0 mg/kg p < 0.0001, mean difference = -3.02, 95% CI: [-4.15, -1.90]. **(H)** As for (G), but with Beam 2. ANCOVA, F(3, 27) = 31.23, p(dose) < 0.0001; Tukey's post-hoc test: 1mg/kg vs 0 mg/kg p = 0.0001, mean difference = -2.14, 95% CI: 95%: [-3.22, -1.07], 3 mg/kg vs 0 mg/kg p < 0.0001, mean difference = -3.39, 95% CI: [-4.50, -2.27]. **(I)** As for (G), but with Beam 3. ANCOVA, F(3, 27) = 24.65, p(dose) < 0.0001; Tukey's post-hoc test 1mg/kg vs 0 mg/kg p = 0.03, mean difference = -1.27, 95% CI: 95%: [-2.43, -0.11], 3 mg/kg vs 0 mg/kg p < 0.0001, mean difference = -3.48, 95% CI: [-4.63, -2.32]. n=7-8 mice (one animal failed to cross Beam 3). Box plots show median ± 95% CI and mean ± 95% in the point plots. ANCOVA was performed to adjust for body weight, with genotype as the factor of interest. Only significant comparisons with control animals are shown. * p<0.05, ** p<0.01, *** p<0.001.

### S2 Fig.

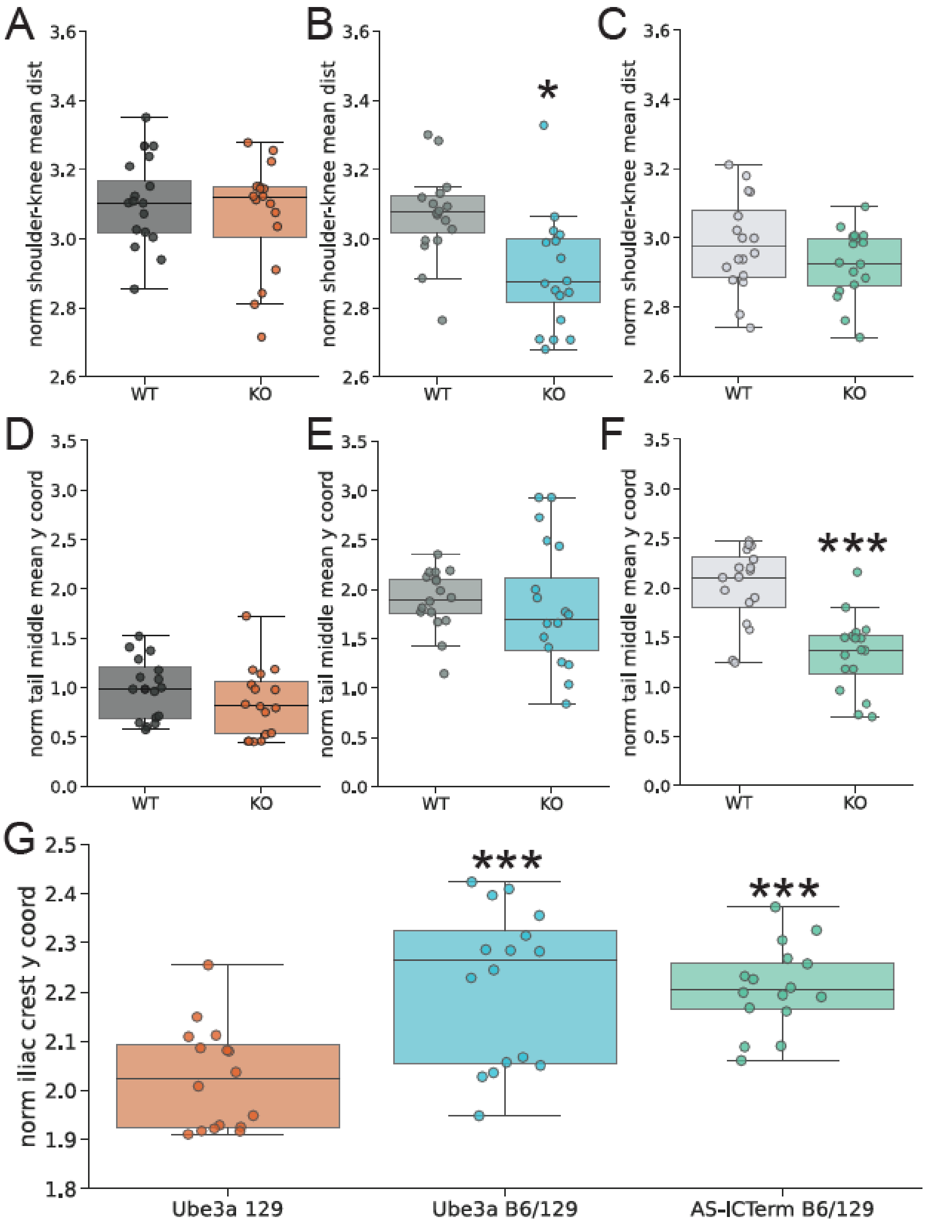


**S2 Fig. Additional comparisons of body posture in mouse models for Angelman Syndrome performing the beam walk. (A)** Shoulder to knee distance in Beam 1 in Ube3a 129 knock out (KO) and wild type (WT) littermates. No significant effect of Genotype: ANCOVA, F(1,29) = 0.12, p = 0.73. **(B)** As panel A, but for Ube3a B6/129 mice. Significant effect of Genotype: ANCOVA, F(1,29) = 5.51, p = 0.03. **(C)** As panel A, but for AS-ICTerm B6/129 mice. No significant effect of Genotype: ANCOVA, F(1,29) = 1.79, p = 0.19. **(D)** Tail middle y-coordinate in Beam 1 in Ube3a 129 knock out (KO) and wild type (WT)

littermates. No significant effect of Genotype: ANCOVA, $F(1,29) = 0.57$, $p = 0.45$. **(E)** As panel D, but for Ube3a B6/129 mice. No significant effect of Genotype: ANCOVA, with weight as a confounding factor, $F(1,29) = 0.13$, $p = 0.72$. **(F)** As panel D, but for AS-ICTerm B6/129 mice. Significant effect of Genotype: ANCOVA, $F(1/29) = 21.20$, $p < 0.0001$. **(G)** Iliac crest y coordinate plotted for KO mice per mouse line. A significant difference was observed between mouse lines (ANCOVA, $F(2,44) = 15.15$, $p < 0.0001$). Tukey's post-hoc test, Ube3a B6/129 KO vs Ube3a 129 KO $p = 0.0002$ (mean difference = -0.19, 95% CI: [-0.29, -0.09]), AS-ICTerm B6/129 KO vs Ube3a 129 KO $p = 0.0002$ (mean difference = -0.18, 95% CI: [-0.29, -0.08]). Sample size = 16 mice per genotype. Box plots show median ± 95% CI. ANCOVA was performed to adjust for body weight, with genotype as the factor of interest. Only significant comparisons with control animals are shown. * $p< 0.05$, *** $p<0.001$.

### S3 Fig

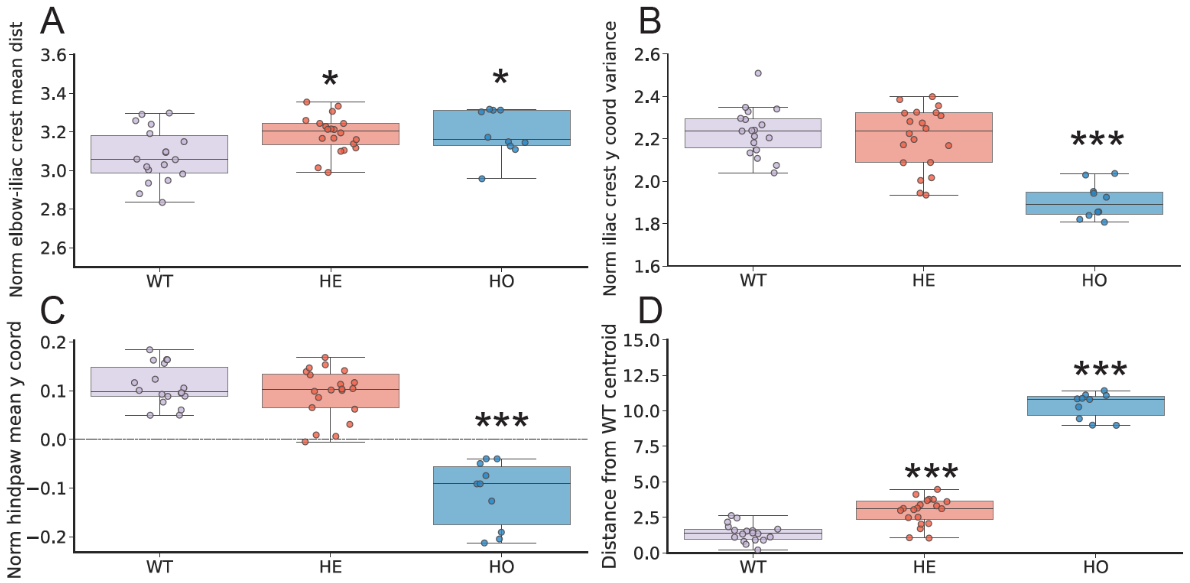


**S3 Fig. Additional analysis of important features discriminating GAT1 mutant mice. (A)** Distance between the elbow and the iliac crest per genotype. Significant effect of Genotype: ANCOVA, F(2, 44) = 4.82, p = 0.01. Tukey's post-hoc test: HE vs WT p = 0.01 (mean difference = -0.11, 95% CI: [-0.21, -0.02]), HO vs WT p = 0.046 (mean difference = -0.11, 95% CI: [-0.23, -0.002]). **(B)** Iliac crest y-coordinate per genotype. Significant effect of Genotype: ANCOVA, F(2, 44) = 12.97, p < 0.0001, and Weight: ANCOVA, F(1, 44) = 26.50, p < 0.0001. Tukey's post-hoc test: HO vs WT p < 0.0001 (mean difference = -0.33, 95% CI: [-0.45, -0.21]), HE vs HO p < 0.0001 (mean difference = 0.30, 95% CI: [0.18, 0.41]. **(C)** Hindpaw y-coordinate per genotype. Significant effect of Genotype: ANCOVA, F(2, 44) = 50.63, p < 0.0001. Tukey's post-hoc test: HO vs WT p < 0.0001 (mean difference = 0.22, 95% CI: [0.17, 0.27]), HE vs HO p < 0.0001 (mean difference = -0.21, 95% CI: [-0.25, -0.16]. **(D)** Euclidean distance from the centroid of the GAT1 WT cluster identified in the space of the first two discriminants following LDA analysis on Beam 1. Significant effect of Genotype (ANCOVA, F(2, 44) = 317.90, p < 0.0001). Tukey's post-hoc test: HO vs WT p < 0.0001 (mean difference = -9.00, 95% CI: [-9.79, -8.21]), HE vs WT p < 0.0001 (mean difference = -1.52, 95% CI: [-2.18, -0.87]). Sample size = 18 WT, 20 HE, 10 HO mice. Plots show median ± 95% CI. ANCOVA was performed to adjust for body weight, with genotype as the factor of interest. Only significant comparisons with control animals are shown. * p< 0.05, *** p<0.001.

### Supplementary files S1 - S7

Supplementary files are available here: https://doi.org/10.5281/zenodo.11074826

**S1 File. 43 pose estimation files used to train foot slip detection.**

**S2 File: List of all 395 features and 50 Prioritized feature lists from across experiments.**

**S3 File. 18 pose estimation files used to verify threshold-methods for foot slip detection.**

**S4 File. Example video from beam walk experiment in mice treated with vehicle, or different doses of diazepam.** Note the change in tail position with increasing doses of diazepam, which was identified as a key feature by Forestwalk that discriminates among treatment groups (see main **Fig. 1**)

**S5 File. Example videos from beam walk experiments in transgenic mice for Angelman syndrome, with comparisons to wild-type control littermates.** For Ub3a 129 KO mice note the more erratic walking style vs. WT controls. For Ube3a B6/129 KO mice, note the more compressed body posture vs. WT controls. For AS-ICTerm B6/129 KO mice, note the change in tail position vs. WT controls (see main **Fig. 3**).

**S6 File. Example video from beam walk experiments in Cohort 1 and Cohort 2 of AS-ICTerm mice.** Note the common postural changes in KO mice vs. WT evident in both cohorts, in particular the lower tail position that is captured by ForestWalk (see main **Fig. 4**).

**S7 File. Example video from beam walk experiments in GAT1 mice.** Note the profound differences in beam walk performance between HO mice vs. WT controls. Differences between HE mice and WT mice are less easily discernible to the human-observer but can be revealed by Forestwalk (see main **Fig. 5**).
